# Neurophysiological signatures in the retrieval of individual autobiographical memories of real-life episodic events

**DOI:** 10.1101/2020.04.13.039024

**Authors:** Berta Nicolás, Xiongbo Wu, Mariella Dimiccolli, Joanna Sierpowska, Cristina Saiz-Masvidal, Carles Soriano-Mas, Petia Radeva, Lluís Fuentemilla

## Abstract

Autobiographical memory (AM) refers to recollected events that belong to an individual’s past. In a classical episodic retrieval experiment in a laboratory, the events to be remembered are words or pictures that have hardly any personal relevance. While such stimuli provide necessary experimental and controlled conditions helping to advance in the understanding of memory, they do not capture the whole complexity of real-world stimuli. Recently, the incorporation of wearable cameras has allowed us to study the cognitive and neural bases of AM retrieval without active participant involvement, and they have been demonstrated to elicit a strong sense of first-person episodic recollection enhancing ecological validity. Here, we provide a new approach to understanding the retrieval of personal events, implementing a convolution network-based algorithm for the selection of the stimuli while monitoring participants’ memory retrieval with scalp EEG recordings over three periods of time after encoding (1 week, 2 weeks, and 6 to 12 months). We also examined an individual with a condition termed Aphantasia that provided more insights into the sensitivity of our protocol in the investigation of individual AM using real-life sequences.

## Introduction

Autobiographical memories (AMs) are specific individualized compilations of our personal past daily life episodic experience. The ability to recollect detailed information about past autobiographical events is a hallmark of episodic memory (Tulving, 2002). However, the vast majority of behavioral and neuroimaging studies of episodic retrieval have used laboratory encoded stimuli, such as words or pictures, as memory probes. While such stimuli provide researchers with tight experimental control over the perceptual qualities, exposure duration, and retention interval of the events being tested, laboratory stimuli lack the richness of most real-world experiences (Chow & Rissman, 2017; Chow et al., 2018; Diamond & Levine, 2020; Nielson et al., 2015; St. Jacques et al., 2011). Thus, it is not unsurprising that performance on standard laboratory-based memory tasks may be largely unrelated to one’s autobiographical retrieval abilities, as demonstrated by individuals with “highly superior autobiographical memory” (LePort et al., 2012; LePort et al., 2017; Patihis et al., 2013) and “severely deficient autobiographical memory” (Fuentemilla, et al., 2018a; Palombo et al., 2015).

A hallmark in the advance of our understating of how episodic memory serves to retrieve real-life autobiographical experiences would be to have methods that allowed the automatic recording of daily life episodes prospectively at the individual level. Through such an approach, researchers would have the opportunity to examine an exhaustive collection of realistic real-life experience material of an individual ahead of sampling control during encoding. Previous research efforts proved effective in cueing AMs sampled at the individual level. Most notably, the use of self-recorded audiotapes or videos documenting selected real-life event experiences has helped characterize the involvement of a core brain network supporting the retrieval of AMs including the medial temporal lobe and the frontal and parietal regions (Levine et al., 2004; Svoboda & Levine, 2009), coordinated via neural oscillatory mechanisms in the range of the theta band (4-8Hz) (Fuentemilla et al., 2018b). However, this approach requires individuals to actively record selected experiences during their daily life routine, and therefore the effectiveness of the retrieval cues may still be partially explained by additional processes engaged during encoding such as selection, organization, and rehearsal of the recorded material.

The recent incorporation of portable technology, such as wearable cameras, to study the cognitive and neural basis of AM retrieval appeared to be a promising venue for addressing the previous concern. This technology allows each individual to capture automatically (e.g., every 30 seconds) face-front sequence of pictures of daily life activity without the need for the participant to be actively engaged in the recording process. Though this is still in its infancy, researchers have already shown that the presentation of pictures acquired with a wearable camera engaged the core AM retrieval network (Cabeza et al., 2004; Rissman et al., 2016) and elicited a strong sense of first-person retrieval in participants, even when they were confronted with others’ pictures depicting the same content (St. Jacques et al., 2011). It proved suitable to test how the hippocampus encodes spatial and temporal properties of our real-life experiences (Nielson et al., 2015). However, given the unfolding nature of our experience, the continuous and automatic photographic process ends up yielding a large number of pictures that require researchers to review and organize them so that they can manually select those, among the large set, that will likely cue a strong sense of episodic recollection in each participant. The fact that only a subsample of pictures are used as retrieval cues in a memory test, and that they are selected based on the experimenter’s criteria, challenges the notion that this approach, as it stands, may be suitable to address fundamental questions of episodic memory, such as “how do we remember or forget single real-life episodes from our past?” and “how do we retrieve individual event episodes over time?”

In the current study, we sought to overcome these issues by recording electroencephalographic activity (EEG) while healthy participants retrieved their own AMs cued by pictures taken automatically (i.e., every 30 seconds) by a wearable camera during one week of daily life routine. To ensure pictures presented during the test cued most of the individual experiences that unfolded during the encoding week, we implemented a convolutional network-based algorithm (SR-clustering, Dimiccoli et al., 2015) on the entire recorded picture set that automatically grouped together temporally adjacent images sharing contextual and semantic attributes, akin to how we conceive what underlies an event episode from a perception and memory perspective (Zacks & Swallow, 2007), which has been recently shown to be a fruitful strategy to catalogue ecologically valid episodic memories (Jeunehomme & D’Argembeau, 2018, 2019). In doing so, the large picture set collected reflecting an entire day’s life activity (e.g., ~400 pictures) is grouped into a workable number of picture subsets (e.g., ~20) depicting sequences of temporally adjacent episodic events (e.g., breakfast at home, commuting to work, meeting with office colleagues, lunch, etc.). Thus, we reasoned that by picking a representative picture from each of the subsets, it would then be possible to investigate whether an individual is capable of recollecting detailed information about the entire set of past experienced events. To assess how the passing of time impacted on the retrieval process of an event episode, we asked each participant to recollect their episodes one week, two weeks, and 6 to 12 months after the encoding period. Additionally, participants were asked to enroll in a separate study that required them to encode and retrieve, one week after the fact, pictures depicting indoor and outdoor scenes. This task, akin to standard lab-based experimental scenarios commonly used in memory research, was thought to help delineate differences between retrieval processes for when participants’ memory was cued by real-life autobiographical vs. lab-based event experiences.

Complementing the behavioral data, we also aimed to analyze well-known neural response activity widely studied in the context of episodic memory retrieval, such as Event-Related Potentials (ERPs) Old/New effects (Friedman & Johnson, 2000) and neural oscillations in the theta range (4-8Hz) (Nyhus & Curran, 2010). More concretely, we expected to identify cued-locked distributed ERP positive responses associated with the successful retrieval of real-life episodic events (Wilding, 1999) and cue-locked modulation of EEG oscillatory activity specifically of the theta band upon successful retrieval. These two neural indexes may well show a distinctive pattern of response as a function of passing of time and lab-based vs. real-life contexts, thereby helping refine the neural underpinnings that underly the retrieval of recent vs. remote autobiographical memories for routine everyday life experiences.

Finally, AM for everyday life activity was examined in an identified a person with confirmed Aphantasia (Zeman et al., 2015; Zeman et al., 2010) using the same wearable camera experimental protocol described previously. GB, as it has been previously described in similar examples (Keogh & Pearson, 2018), was an individual that showed a specific lack of visual imagination while preserving the rest of the cognitive abilities intact. Visual imagination is critical in determining the retrieval of high-quality episodic memories (Greenberg & Rubin, 2003) and it has been suggested as a feature to help distinguish individuals in terms of their ability to recollect AMs (Palombo et al., 2018). Here, we aimed to use our novel wearable camera-based experimental design to characterize GB’s AM and with that to gain insight into the sensitivity of our protocol to scrutinize AM at the single subject level.

## Material and methods

### Participants

Sixteen healthy participants (8 females) participated in experiments 1 and 2. The range in age was 22-37 years old (average age=27.68 years old (SD=4.22)). All participants provided informed written consent for the protocol approved by the Ethics Committee of the University of Barcelona. Participants received financial compensation for their participation. Two of the participants could not complete the follow-up in experiment 1 (see details below) and these participants were excluded from all analyses.

In addition, we examined a person with Aphantasia, referred to as case GB (not actual initials). GB was a 40-year-old male who contacted our research group after reading an article about Aphantasia in popular media. He recognized his personal experience with the information described in the article. The Vividness of Visual Imagery Questionnaire (VVIQ) (Marks, 1973) confirmed that GB had Aphantasia as he scored 8 (z = – 5.72) on a scale of 0 (minimum or complete lack) to 224 (maximum) in the test. GB reported no neurological, ophthalmological, or psychiatric etiology of his reported lack of imagery. Neuropsychological results are described in supplementary Table 1. Structural MRI (see Methods for details) showed minor white matter high intensities and borderline fronto-temporal atrophy, both within the normal limits for his age. GB’s MRI scan was compared to a control sample of six participants (males, average age = 30.83 years old (SD = 3.87), average education years = 14.50 (SD = 4.37)).

## Experiment 1: Retrieval of real-life memories

### Design overview

Participants were asked to come to the training session a few days before starting the study. They were informed about how the camera worked and they read, understood, and signed the informed consent. We took special care in providing details on privacy issues so that all participants were fully aware of them before the study began. In the current study, participants wore the wearable camera for a period of 5-7 consecutive days (mean = 6.64 days, SD = 0.63), including both week and weekend days. The data collection period was established from morning to evening (between 12-14 hours per day). Once participants finished data collection, they were requested to return the materials to be processed. Participants confirmed not having checked the pictures during the recording week. Participants returned to the lab to be tested one week later (T1), two weeks later (T2), and from 6 to 14 months after the last day of data collection (Follow-up, hereafter FU; mean = 10.87 months, SD = 2.02 months). See Figure 1 for a summary of the experimental design.

**Figure 1.**
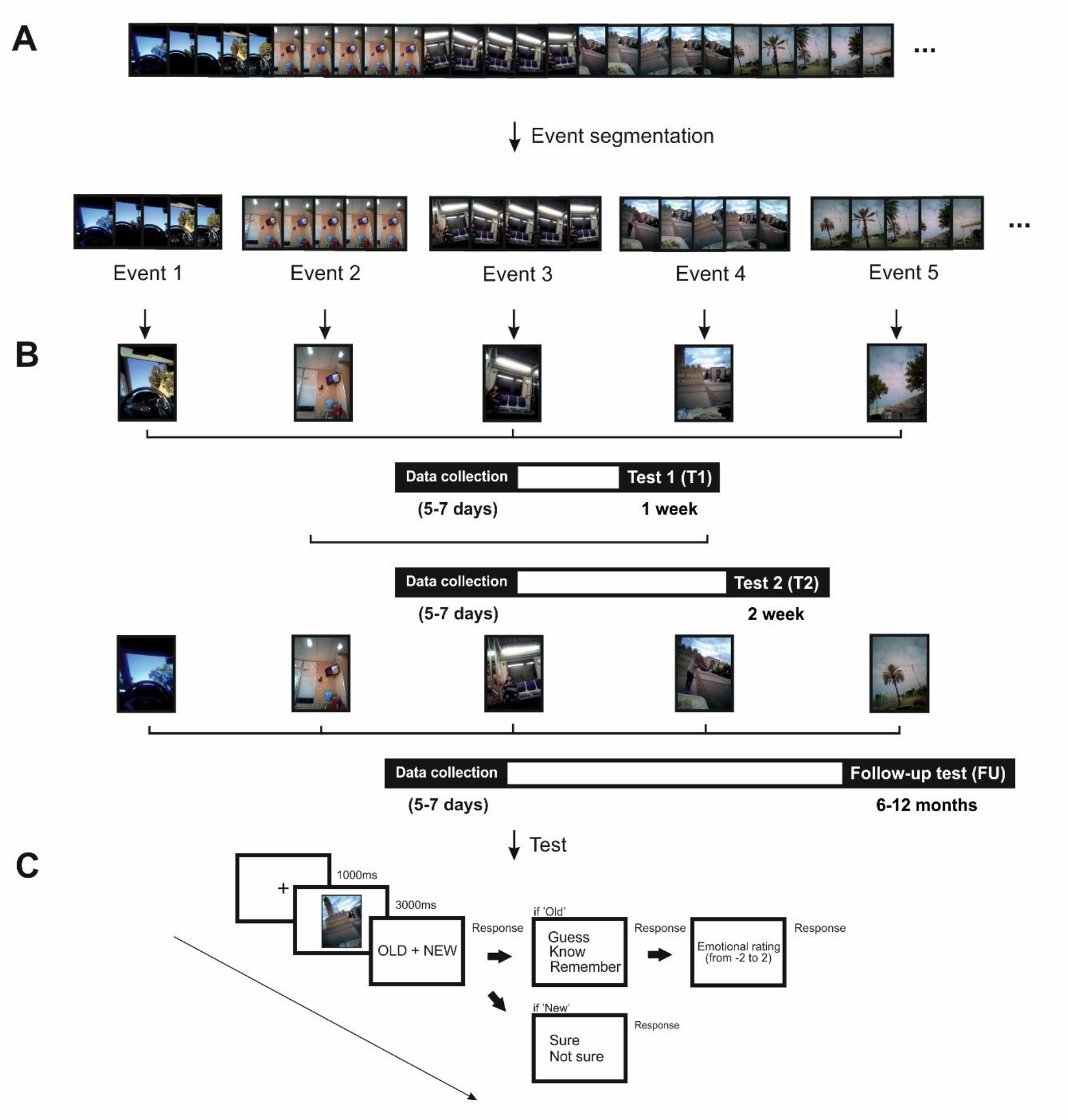
**(A) Event segmentation approach**. Example of a stream of pictures obtained from one participant after one week of data collection. The implementation of the SR-Clustering algorithm allows the automatic organization of picture sequences into a set of meaningful events by the identification of similar context and semantic features. **(B) Experimental design.** One representative photo from each event sequence of pictures was selected and distributed to each of the three memory tests that followed the picture collection week. A recognition memory for the was implemented one week (T1), two weeks (T2) and from 6 to 12 months (Followup test (FU)) after the data collection. Events extracted from picture sequences were numbered consecutively and pictures related to even numbered events were used as memory cues in T1 test and odd numbered event pictures in T2 test. Different pictures from same events tested in T1 and T2 were selected as cues in FU test. **(C) Recognition memory task**. Pictures were presented on the screen for 3000 ms. Afterward, an ‘Old/New’ question appeared on the screen. If the pictures were seen as ‘Old’, a “Remember/Know/Guess” appeared on the screen. Next, participants were asked to rate their emotional response towards the event indicating whether was positive or negative on a scale ranged from 2 (maximum positive) to −2 (maximum negative) being 0 neutral emotion. If the picture was seen as ‘New’, ‘Sure/Not sure’ judgment appeared on the screen.

### Wearable camera

We used the wearable Narrative clip 2 camera^®^ (http://getnarrative.com/) with a camera sensor of 8MP and a resolution of 3264×2448(4:3). The camera was programmed to automatically take images every thirty seconds and produced pictures with an egocentric viewpoint. Participants were instructed to wear the device on a lanyard around the neck. Narrative clip 2 incorporated a downloading app that allowed participants to download the pictures directly to a hard drive. Participants were instructed to not watch the pictures until the experiment finalized. None of the participants reported having done so at the end of the study.

### Picture selection

We implemented a deep neural network-based algorithm, SR-Clustering, to automatically organize the stream of each participant’s pictures into a set of temporally evolving meaningful events (Dimiccoli et al., 2015). The algorithm segments picture sequences into discrete events (event segmentation; e.g., having breakfast in a kitchen, commuting to work, being in a meeting) on the basis of its ability to identify similar contextual and semantic features from the picture stream. See Figure 1A for the event-segmentation approach.

The implementation of the SR-Clustering algorithm provided a variable number of discrete events for each participant per day, and each event included 8 to 20 pictures. Each participant’s events were then manually inspected and those which displayed non-meaningful episodes (e.g., all pictures were blurred, or when the camera was pointing to the roof or was blocked by clothes) were discarded from the study. Picture events that described interactions with participants’ relatives were excluded from the study. Three independent experimenters rated and selected the set of event pictures for each participant on the basis of these criteria, and only those events that were consistently selected by the three raters were included in the final set of picture events in the study (average number of events across participants Mean = 322, SD = 127). Note that the consistency across experimenters was set to ensure that the events captured by the algorithm were meaningful and did not involve implementing a subjective inclusion/exclusion selection criterion to which events should be included later in the memory test by the experimenters. The variability observed in the number of events between participants reflected the diversity of each participant’s daily life activities (e.g., a person working indoors for 8 hours results in fewer events compared with people working outdoors).

Once the images were organized into discrete events, we selected a representative picture from each event, thereby ensuring most of the past episodic experience was brought into the test. We then numbered the sequence of event pictures and assigned even-numbered pictures to be used as memory cues for test T1 and odd-numbered pictures to be included in the T2 test. FU included picture cues used in T1 and T2 in the same proportion. Pictures cues presented to one participant depicting her own past (Old) were also presented to another participant as New images. This ensured that differences between Old and New pictures presented to each participant were only based on the image’s direct link to ones’ past while preserving the rest of the characteristics intact during the test (e.g., angle of view, picture image features, description of routine daily life activities). See Figure 1B for a summary of the experimental design.

By design, none of the participants were friends with each other, and we never encountered an instance where two concurrently enrolled participants came into direct contact with one another while wearing their cameras.

### Recognition memory task

In the test, pictures were presented on the screen for 3000 ms. Afterwards, when an “Old/New” question appeared on the screen, participants were required to judge whether the picture reflected an event from the participant’s own daily life (Old) or was experimentally novel, signaling with the right index and middle fingers, respectively. Next, participants were asked to judge whether they were “Sure/Not sure” when indicated an image was “New” and “Remember/Know/Guess” when images were seen as “Old”. Participants were instructed that “Guess” referred to when they had no contextual memory reference for what was depicted in the image, but they recognized the content as being from their own life (e.g., viewing ones’ living room). “Know” was the signal for when the visual content in the picture was highly familiar but the subject could not determine what unfolded in it, perhaps because the event on the test was part of a routine (e.g.: playing football on Thursdays), while “Remember” was the signal for when the picture elicited a vivid memory of that specific event and it could be located in time. Participants’ ability to order each event depicted in the pictures along the encoding week was tested afterward more concretely, when they were asked to indicate whether the pictures depicted an event that took place at the “beginning, middle or end” of the encoding week as the image appeared on the screen. Finally, participants were asked to rate the degree to which each of the pictures elicited an emotional response and to indicate whether it was positive or negative on a scale that ranged from 2 (maximum positive) to −2 (maximum negative), with 0 being neutral emotionally. See Figure 1C for a summary of the recognition memory task. Temporal ratings are not shown in the design overview.

## Experiment 2: Retrieval of lab-based memories

The experimental design was similar to experiment 1, but differed mainly in that images depicting real-life experiences were replaced by neutral images of indoor and outdoor scenes extracted from previous experiments (e.g., Bunzeck & Düzel, 2006; Fuentemilla et al., 2010). Thus, the design involved an encoding phase and a test phase administered after each participant finished the retrieval session from the FU condition in experiment 1.

In the encoding phase, participants were instructed to indicate whether scene pictures were indoor or outdoor images. There were 80 scenes (40 indoor and 40 outdoor, presented in random order). Each scene was presented for 2000 ms preceded by a 1500 ms fixation period and followed by the text “indoor/outdoor” that prompted participants’ response (responding with the index or middle finger of their right hand). A period of 10 minutes of rest separated the study from the test phase.

In the test phase, a scene picture was presented on the screen for 3000 ms. Afterwards, when an “Old/New” question appeared on the screen, participants were required to judge whether the word was presented in the previous study phase (Old) or was experimentally novel (New) with the right index and middle finger, respectively. The test phase included 160 scene images in total (80 Old and 80 New, randomly presented). Thereafter, as in experiment 1, a confidence judgment task followed. Here, new judgments were followed by “Sure/Not sure” and old judgments were followed by “Remember/Know/Guess”. Participants were instructed to make confidence judgments following old judgments with respect to their ability to vividly retrieve the contextually associated information related to the image during encoding. They were instructed to respond “Guess” when they were unsure about their previous Old judgment, “Know” when they recognized the scene image but could not retrieve any contextual feature linked to it, and “Remember” when the scene image brought a vivid recollection of the specific context that surrounded the encoding of that particular image during encoding.

### EEG recordings and preprocessing

EEG was recorded (band-pass filter: 0.01–250 Hz, notch filter a 50Hz, and 500 Hz sampling rate) from the scalp using a BrainAmp amplifier and tin electrodes mounted in an electrocap (Electro-Cap International) located at 29 standard positions (Fp1/2, Fz, F7/8, F3/4, FCz, FC1/2, FC5/6, Cz, C3/4, T3/4, Cp1/2, Cp5/6, Pz, P3/4, T5/6, PO1/2, Oz) and at the left and right mastoids. An electrode placed at the lateral outer canthus of the right eye served as an online reference. EEG was re-referenced offline to the linked mastoids. Vertical eye movements were monitored with an electrode at the infraorbital ridge of the right eye (EOG channel). Electrode impedances were kept below 3 kΩ. EEG was band-pass filtered offline at 0.1 – 40Hz. Independent Component Analysis (Delorme & Makeig, 2004) was applied to the continuous EEG data to remove blinks and eye movement artefacts. EEG data from two participants were lost due to technical problems and were not able to be included in the rest of the EEG analysis.

### Event-related potentials (ERPs) analysis

The continuous sample EEG data were then epoched into 3100 ms segments (0 to 3000 ms relative to trial onset), and the pre-stimulus interval (−100 to 0 ms) was used as the baseline for baseline removal procedure. Trials exceeding ± 100 μV in EEG and/or EOG channels within −100 to 3000 ms time window from stimulus onset were rejected offline and not used in ERPs and time-frequency analysis (see details below). For each participant, we obtained trial epochs that were separately catalogued as belonging to Old and New conditions.

### Time-frequency analysis

The power of neural oscillatory activity was calculated by means of the continuous complex Morlet wavelet (Grossmann & Morlet, 1984). It is a biologically plausible wavelet modulated by a Gaussian function which depends on the number of cycles the sinusoidal wave segment comprises. In the current study, the cycles of the Morlet wavelets used for convolution ranged from 4 to 10, increasing logarithmically as frequency increased. We adopted this modified wavelet approach to optimize the trade-off between the temporal resolution at lower frequency band and the frequency resolution at the higher frequency band. For all conditions in experiment 1 and 2, time-frequency analysis was carried out on a single trial basis, with epochs of 3500 ms time-locked to the presentation of photo starting at 500 ms before its onset. The convolution with Morlet wavelet was conducted for each frequency value from 1 Hz to 40 Hz, with 50 steps increasing logarithmically. Power values for each frequency were averaged across trials for each channel and then baseline-corrected by decibel conversion.

### Cluster-based statistics of the ERP and Time-frequency data

To account for ERP differences elicited by Old and New pictures a cluster-based permutation test was used (Maris & Oostenveld, 2007) to identify clusters of significant points in the resulting spatiotemporal 2D matrix (time and electrodes) in a data-driven manner and addressing the multiple-comparison problem by employing a nonparametric statistical method based on cluster-level randomization testing to control for the family-wise error rate. Statistics were computed for each time point, and the spatiotemporal points whose statistical values were larger than a threshold (p < 0.05, two-tail) were selected and clustered into connected sets on the basis of x, y adjacency in the 2D matrix. The observed cluster-level statistics were calculated by taking the sum of the statistical values within a cluster. Then condition labels were permuted 1000 times to simulate the null hypothesis, and the maximum cluster statistic was chosen to construct a distribution of the cluster-level statistics under the null hypothesis. The nonparametric statistical test was obtained by calculating the proportion of randomized test statistics that exceeded the observed cluster-level statistics.

To assess for differences between Old and New conditions at the time-frequency level a similar statistical approach was adopted. However, clusters (p < 0.05, two-tail) were determined by connected sets of data samples that were contiguous on the basis of temporal, frequency, or spatial adjacency in the 3D matrix. Cluster statistics and null distribution were created following the same approach as for the ERP statistical approach.

## Aphantasia: A case study

GB ability to retrieve individual real-life memories was examined by implementing the experimental design described in Experiment 1. More specifically, this person carried the wearable camera for 7 consecutive days and returned to the lab to perform a retrieval task while scalp EEG was acquired 1 week after encoding. The selection of the pictures depicting GB’s past events (Old) and those depicting events from others (New) followed the procedure indicated in experiment 1 (total number of events = 199). EEG recording, ERP, and timefrequency analysis were similar to those described previously.

Additionally, an extensive neuropsychological screening was implemented for GB (results included in supplementary Table 1) as well as structural Magnetic Resonance Imaging (MRI) scanning session. GB was scanned in a 3T Philips Ingenia MRI Scan (Philips Medical System) equipped with a 32-channel head coil, at the Hospital Universitari de Bellvitge of Barcelona. We used a three-dimensional fast-spoiled gradient, inversion-recovery sequence with 233 contiguous slices (repetition time, 10.43 msec; echo time, 4.8 msec; flip angle, 8°) in a 24-cm field of view, with a 320 × 320-pixel matrix and isotropic voxel sizes of 0.75 × 0.75 x 0.75 mm.

EEG and MRI data were compared to those for the group by means of the Crawford *t* statistic, which is especially suited to assessing comparisons between single subjects and group sample data (Crawford & Howell, 1998).

## Results

### Experiment 1

#### Behavioral results

Participants were highly accurate in correctly distinguishing pictures that depicted their own past (Old pictures) from those that belonged to others’ past (New pictures) (Figure 2A). False Alarms (T1 test: Mean (M) = 0.05, Standard Deviation (SD) = 0.05; T2 test: M = 0.03, SD = 0.03; FU test: M = 0.03, SD = 0.02) and Omissions (T1 test: M = 0.08, SD = 0.06; T2 test: M = 0.09, SD = 0.06; FU test: M = 0.11, SD = 0.09) were very rare in all tests. However, a repeated measures ANOVA, including the three-memory test conditions (T1, T2, and FU) as a within-subject factor in the analysis, revealed that hit rate differed significantly across them (F (2,26) = 4.94, p = 0.01) (Figure 2A). A series of paired t-test comparisons showed that hit rate decreased as a function of time from encoding. Thus, significant differences were obtained when T1 and FU hit rate were compared (t(13) = −2.68, p = 0.02) but not for T1 and T2 (t(13) = 1.37, p = 0.19), nor T2 and FU (t(13) = 1.93, p = 0.07). These differences cannot be accounted for by a general decrease in performance over time as participants’ ability to identify New images (i.e., correct rejections) was similar in the three tests (F (1.29, 16.85) = 2.14, p = 0.16). Correct rejections: T1 test: M = 0.94, SD = 0.05; T2 test: M = 0.97, SD = 0.03; FU test: M = 0.97, SD = 0.02).

**Figure 2.**
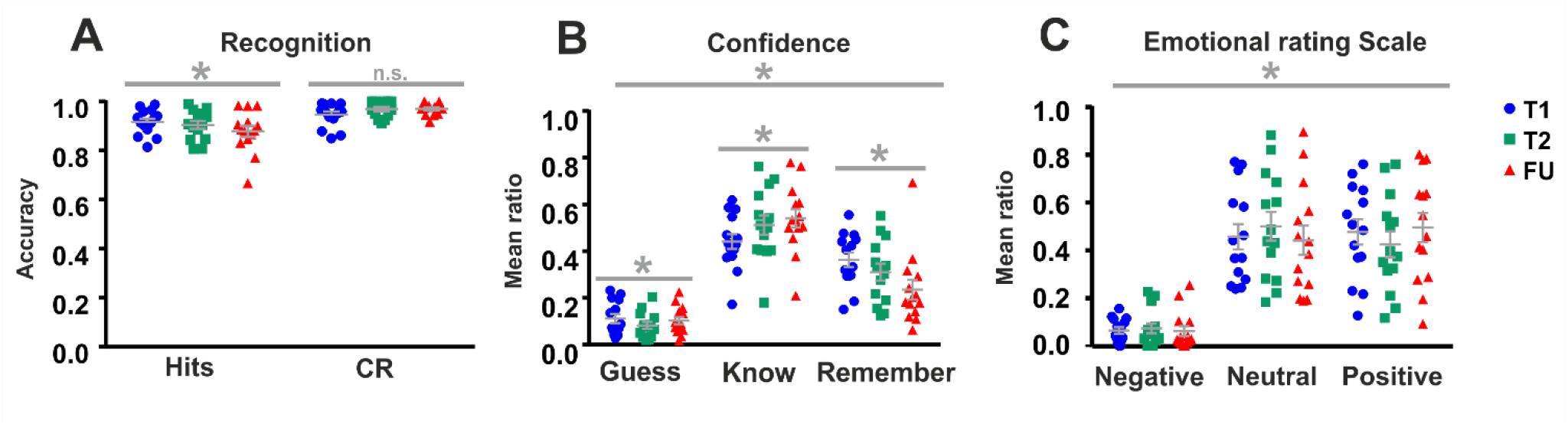
Behavioral data in healthy young participants from Experiment 1. **(A)** Averaged participants’ accuracy and correct rejections in selecting their own events (old) compared to others’ events (new) over three time periods (T1: one week, T2: two weeks, FU: 6 to 12 months). **(B)** Confidence judgments over three time periods (T1, T2, and FU). **(C)** Emotional ratings over three time periods (T1, T2, and FU). The asterisks indicate significant (p < 0.05) effects. Error bars represent SEM.

A repeated measures ANOVA including participants’ confidence ratings (Guess, Know, and Remember) on each of the three memory tests (T1, T2 and FU) revealed that participants distribution of responses concentrated around specific ratings (effect of rating: F(2,26) = 4.94, p = 0.01), and that they fluctuated differently as a function of time (effect of test: F(2,26) = 37.99, p < 0.01; interaction rating x test: F(1.88,24.42) = 5.51, p < 0.01) (Figure 2B). To disambiguate this effect, we ran separate repeated measures ANOVAs including memory test as a within factor for each of the confidence ratings. We found that differences across tests were in “Know” (F(2,26) = 4.93, p = 0.01) and “Remember” (F(2,26) = 6.31, p < 0.01), but not in the distribution of “Guess” responses (F(2,26) = 2.65, p = 0.09). A series of paired t-tests revealed that “Know” responses increased over memory tests (T1 vs T2: t(13) = −2.93, p = 0.01; T2 vs FU: t(13) = −0.74, p = 0.47; T1 vs FU: t(13) = −3.02, p = 0.01) and that “Remember” responses were mostly located in memory test T1 (T1 vs T2: t(13) = 2.03, p = 0.06; T2 vs FU: t(13) = 1.74, p = 0.11; T1 vs FU: t (13) = 3.43, p < 0.01).

We next examined participants’ proportion of emotional rating to the correctly retrieved pictures over tests (T1, T2 and FU). This showed that participants rarely indicated maximum negative (i.e., −2 in the scale) or positive (i.e., +2) emotion in the tests (negative: M = 1.14%, SD = 1.94%; positive: M = 14.35%, SD = 13.61%; respectively across the three tests). Therefore, we grouped −2 and −1 responses as negative and +1 and +2 as positive, leaving 0 as indicating neutral emotion. A repeated measures ANOVA, including time (T1, T2, and FU) and emotion (negative, neutral, and positive) as within subject factors, revealed a main effect of emotion (F (1.13,14.65) = 22.52, p < 0.01) but a non-significant emotion x time interaction (F(1.62,21.11) = 0.72, p > 0.05) (Figure 2C). To identify the source of this main effect, we averaged the proportion of each emotion’s category across the three tests. A series of paired t-test comparisons revealed that most pictures elicited neutral emotion in the participants (neutral vs negative: t(13) = 8.02, p < 0.01; neutral vs positive; t(13) = 0.01, p = 0.99) and positive emotion (positive vs negative: t(13) = 7.53, p < 0.01).

Finally, participants’ response accuracy to temporal order memory was at random (M = 0.49, SD = 0.22), thereby indicating this test was not suitable to capture the participants’ ability to retrieve temporal representations of their own AMs.

#### ERPs results

Our analytical strategy was to first assess whether Old and New images elicited different patterns of brain activity in the participants. To address this issue, we averaged, at the participant level, the ERPs elicited by Old and New image conditions across tests and ran a cluster-based permutation test between these two conditions. This analysis revealed a significant cluster showing that Old images elicited higher ERP positive amplitude from 400 ms at stimulus onset, which lasted over the rest of the temporal window in the analysis and comprised a large distributed area over the scalp (Figure 3A). To assess how the identified Old/New ERP effect varied as a function of memory test, we submitted the averaged ERP activity from the cluster to a repeated-measures ANOVA including experimental condition (Old, New) and memory test (T1, T2, and FU) as a within-subject factor. As expected, this analysis revealed a main effect of condition (F(1,11) = 60.24, p < 0.01) and a significant condition x memory test interaction (F(2,22) = 6.45, p < 0.01) (Figure 3B) but not a main effect of memory test (F(2,22) = 0.79, p = 0.47). However, separate repeated-measures ANOVAs for ERP values to Old and New images revealed that none of them showed a statistically significant effect as a function of memory test (Old: F (2,22) = 0.66, p = 0.53; New: F (2,22) = 2.58, p = 0.09).

**Figure 3.**
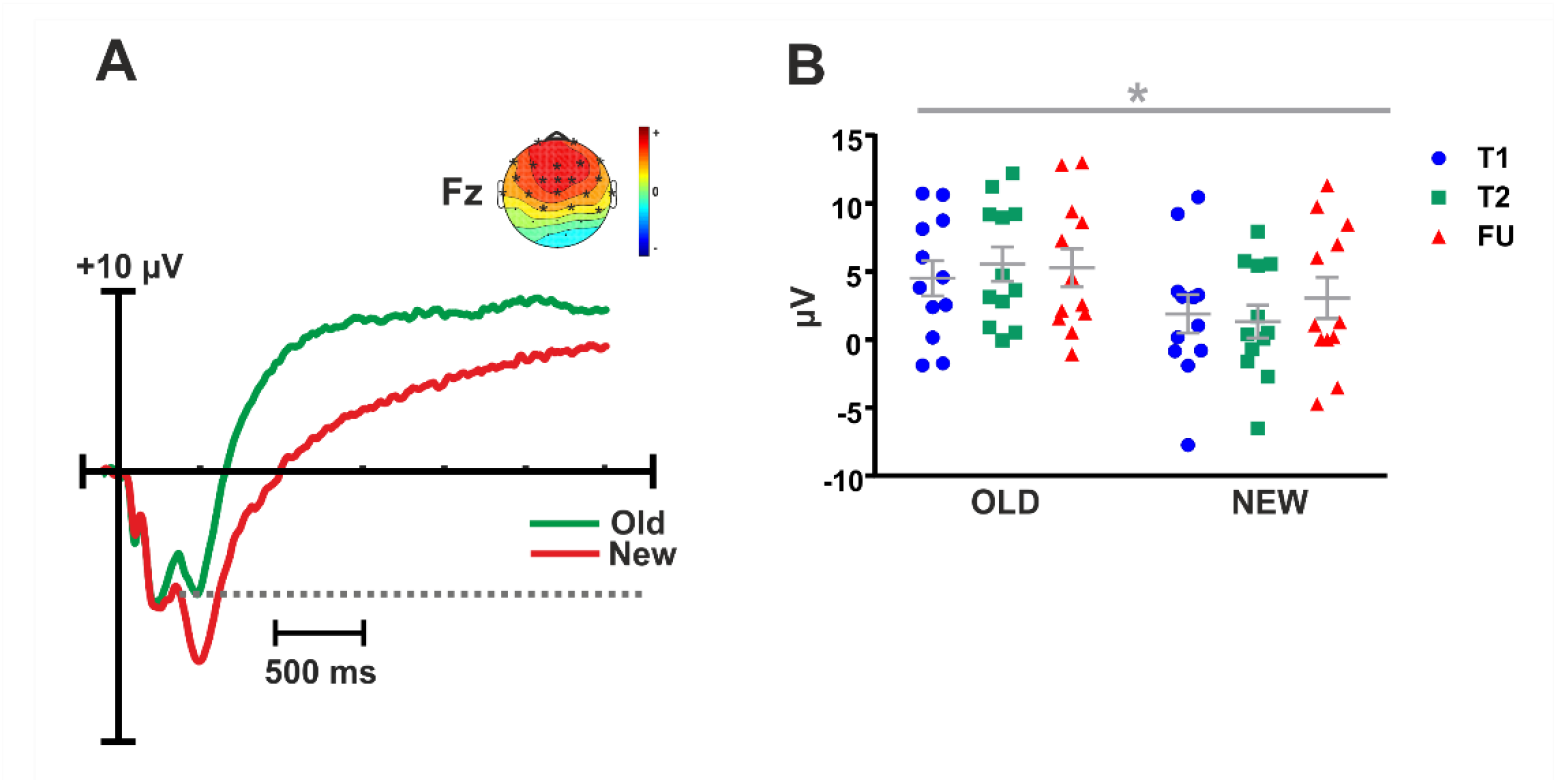
**(A)** Across participants grand-average event-related potentials (ERPs) for Experiment 1 for Old and New conditions at Fz. A cluster-based permutation analysis between the two conditions revealed that Old pictures elicited greater ERPs amplitude than did fronto-central scalp regions. Dashed line indicates the temporal window of significance (p < 0.05, corrected). **(B)** Cluster-averaged individual ERP data for each of the three recognition memory tests and conditions (T1, T2, and FU). * indicates p < 0.05. Error bars represent SEM.

#### Time-frequency results

Following the ERP analytical strategy, we first implemented a cluster-based permutation test to identify, in a data-driven manner, for the existence of a main Old/New difference pattern of neural oscillatory response along the temporal x spatial x spectral dimension. Thus, spectral power measures elicited at the onset of Old and New correct responses were averaged across the three tests (T1, T2, and FU) and were then compared. This analysis revealed the existence of a significant cluster initiating at around 1000 ms from stimulus onset comprising mostly low-frequency activity in the theta range (4-8Hz) (Figure 4A). More specifically, these results showed that Old responses were accompanied by a decrease in theta power and that this effect was over frontal and central regions of the scalp. A repeated-measures ANOVA, including as within-subject factors memory test (T1, T2 and FU) and image type (Old and New), showed a main effect of image type (F(1,11) = 56.85, p < 0.01), no significant main effect of memory test (F(2,22) = 0.33, p = 0.73), but the existence of a significant image type x memory test interaction (F(2,22) = 5.15, p = 0.01) (Figure 4B). These results suggest that theta Old/New effect differed between memory tests. However, paired t-test analysis showed that Old vs New theta differences were significant at T1 (t(11) = 4.21, p < 0.01), at T2 (t(11) = 7.21, p < 0.01), and at FU (t(11) = 3.69, p < 0.01), thereby hindering the possibility of establishing the source of this interaction clearly in our data.

**Figure 4.**
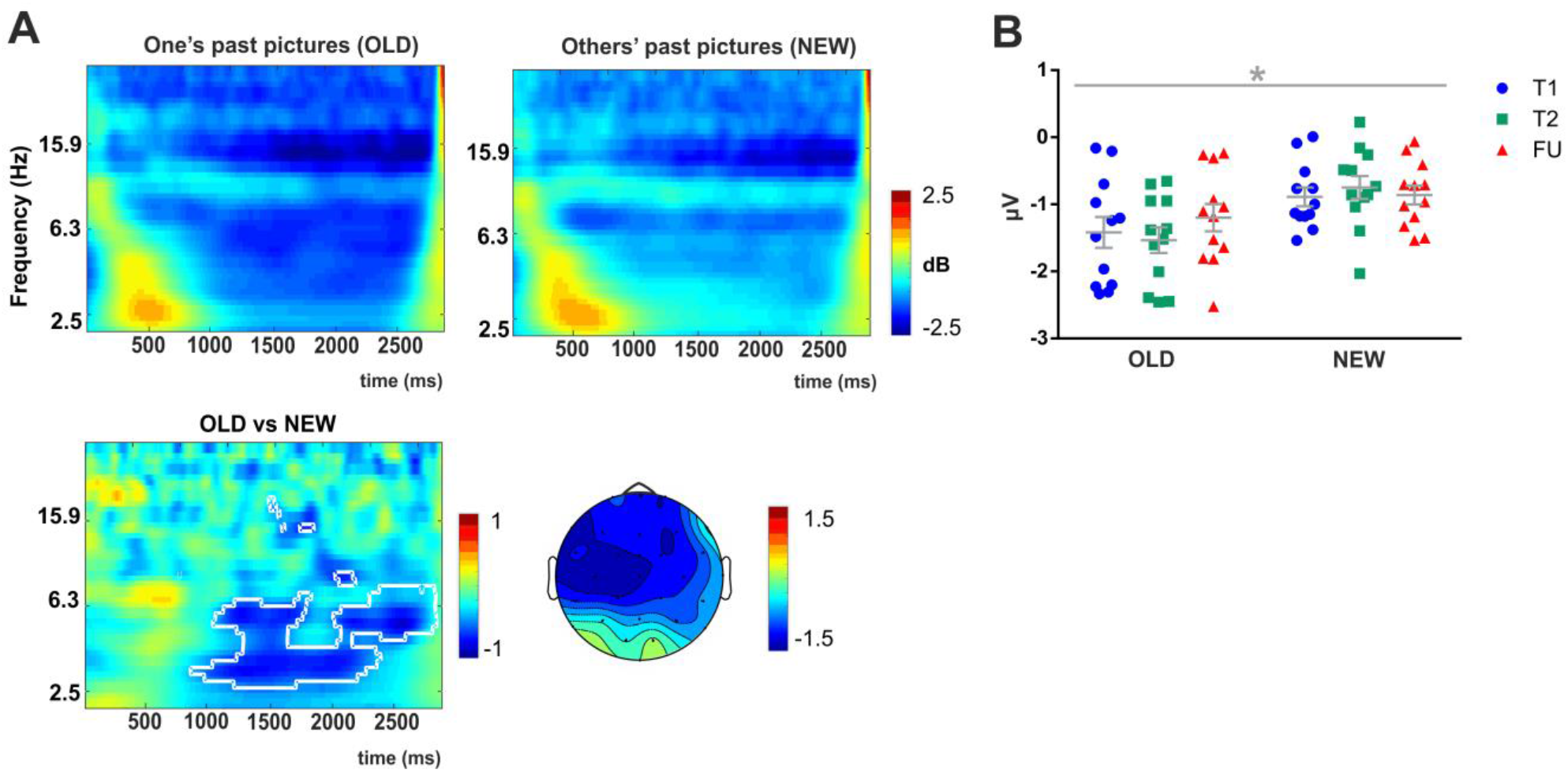
Theta oscillations in Experiment 1. **(A)** Group-averaged changes in spectral power over the three periods of time (averaged over all scalp sensors) elicited by pictures related to own personal events (old) compared to others’ events (new). Decreased theta power was observed in old responses over fronto-central regions of the scalp. **(B)** Average theta power over three periods of time (T1, T2, and FU). Significant (p < 0.05) interaction effect is indicated with a grey star. Error bars represent SEM.

### Experiment 2

#### Behavioral results

Participants’ hits and correct rejection (CR) rates are displayed in Figure 5A. Overall, participants’ performance was much poorer in this experiment. To assess this statistically, we compared participants’ Hits and CRs in experiment 1 and in experiment 2. This analysis used participants’ behavioral data obtained in condition T1 from experiment 1, as this shared a one-week time frame between encoding and retrieval in the two experiments. This analysis confirmed significant differences in the proportion of Hits (F(1,13) = 47.09 p < 0.01) and in CRs (F(1,13) = 31.12, p < 0.01) between experiments, thereby indicating, as expected, that participants were much accurate in recognizing pictures related to their own past real-life experience than those encoded “artificially” in a lab-context.

**Figure 5.**
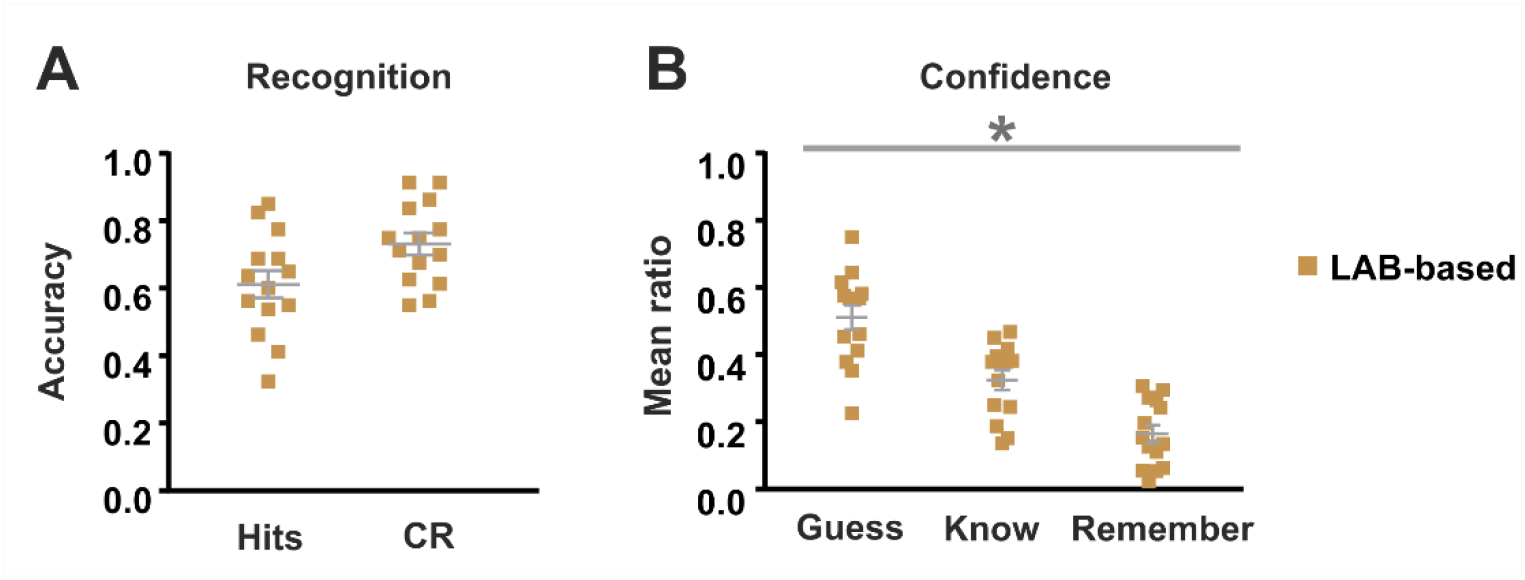
Behavioural data in healthy young participants from Experiment 2. **(A)** Averaged participants’ accuracy and correct rejections in selecting pictures previously encoded (old) compared to pictures never seen (new). **(B)** Confidence judgments based on laboratory stimuli. The asterisks indicate significant (p < 0.05) effects. Error bars represent SEM.

In addition, a repeated-measures ANOVA of participants’ confidence ratings revealed that the ratings following correct Old responses were unequally distributed (F (2,26) = 20.72, p < 0.01; Figure 5B). More specifically, participants tended to rate the correctly recognized scene images as having “guessed” more frequently than “remembered”, thereby indicating that memory accuracy was, overall, quite low. When comparing confidence ratings provided in the analogous experiment 1, we found a significant interaction effect of confidence (F (2,26) = 42.26, p < 0.01), thereby indicating confidence rating distribution differed between experiments. These results indicate that participants’ sense of recollection for laboratory-based material was not as vivid as when they judged their own stimuli.

#### ERPs results

EEG data were analyzed as in experiment 1, by contrasting ERP patterns of activity elicited by Old and New images through a data-driven cluster-based permutation test. This analysis revealed a significant cluster showing that Old images elicited higher ERP positive amplitude at 400 ms from stimulus onset. However, compared to the ERP differences observed in experiment 1, this cluster of EEG activity was much shorter and less well distributed over the scalp (Figure 6A). Nevertheless, a repeated-measures ANOVA including response type (Old and New) and experiment (T1 from experiment 1 and experiment 2) as within-subject factor revealed that the Old/New ERP effects were statistically similar in the two experiments (main effect of experiment: F(1,9) = 1.96, p = 0.19; experiment x response type interaction: F(1,9) = 3.29, p = 0.10).

**Figure 6.**
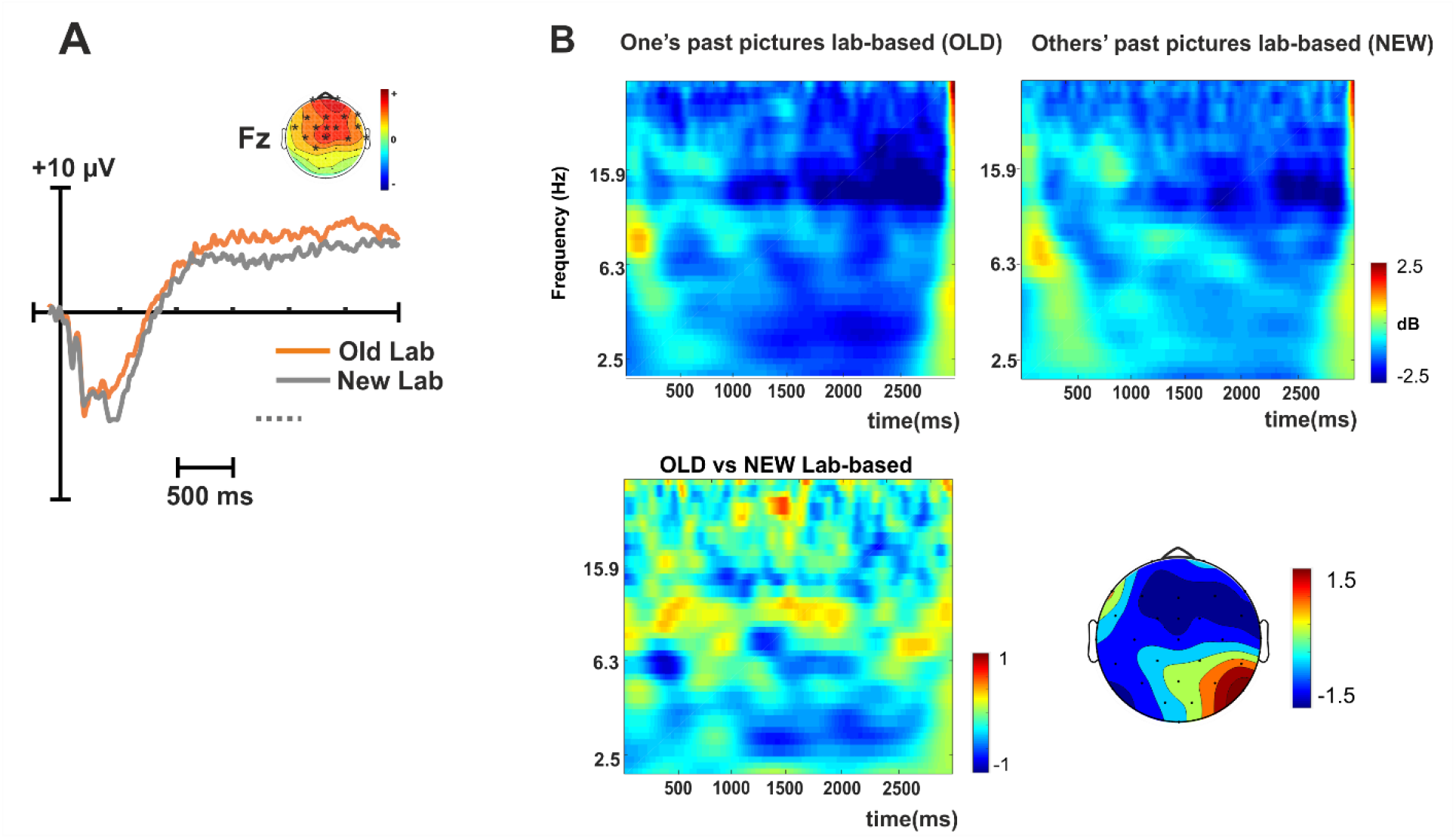
**(A)** Across-participants grand-average event-related potentials (ERPs) for Experiment 2 for Old and New conditions at Fz. A cluster-based permutation analysis between the two conditions revealed that Old pictures elicited greater ERP amplitudes than did fronto-central scalp regions. Dashed line indicates the temporal window of significance (p < 0.05, corrected). **(B)** Group-averaged changes in spectral power (averaged over all scalp sensors) elicited by pictures related to own personal events (old) compared to others’ events (new). Decreased theta power was observed in old responses over fronto-central regions of the scalp.

#### Time-frequency results

The implementation of a cluster-based permutation test assessing differences between Old and New responses to scene images encoded in the laboratory yield non-significant results along the temporal x frequency x spatial (scalp sensors) dimensions (Figure 6B). Thus, contrary to the theta Old/New effects found in experiment 1, differences were not observable in experiment 2. Nevertheless, to assess the extent to which the Old/New theta effects found in experiment 1 differed from the same theta modulations in experiment 2, we ran a repeated-measures ANOVA including response type (Old and New) and experiment (T1 from experiment 1 and experiment 2) as within-subjects factors. Given the null effects found in experiment 2, theta power in this experiment was extracted by averaging data points that were within the temporal x frequency x spatial cluster data points identified in experiment 1. This analysis confirmed the main effect of image type (F(1,9) = 9.57, p = 0.013) but uncovered no significant main effect of the memory test (F(1,9) = 1.96, p = 0.195) nor type x memory test interaction (F(1,9) = 3.29, p = 0.103), which hindered the possibility of concluding that the Old/New ERP effect was greater in one of the experiments. However, note that the clusters of activity included in this analysis were much larger in ERPs from experiment 1 than in ERPs from experiment 2.

### Case GB (Aphantasia)

#### Behavioral data

GB was highly accurate in identifying images that belonged to his past (Hit rate = 0.91) and those that did not (Correct Rejection rate = 0.05). However, hits were mainly “Known” judgments (rate of 0.57; Guess rate = 0.03 and Remember rate = 0.29), thereby indicating that his ability to correctly recognize Old pictures was usually not accompanied by a detailed or vivid recollection of the cued AM event episode.

In order to evaluate statistically the extent to which case GB’s behavioral performance was similar to the sample of participants from experiment 1, we compared each of the behavioral measures between the single case and group sample at T1 by means of the Crawford t-test. However, none of the comparisons proved to be statistically significant (all p > 0.05).

#### EEG (ERPs and Time-Frequency data)

A clear Old/New ERP effect was found in GB (Figure 7A). ERPs to Old and New responses were similar to those found in the sample group that participated in experiment 1. Thus, differences between ERPs were displayed at frontal and central scalp regions and they began at around 400 ms from stimulus onset and persisted over the 3000 ms epoch window. This was confirmed by the finding that the ERP to Old and New stimuli in GB and in the participants from experiment 1 was statistically similar (all, p > 0.05).

**Figure 7.**
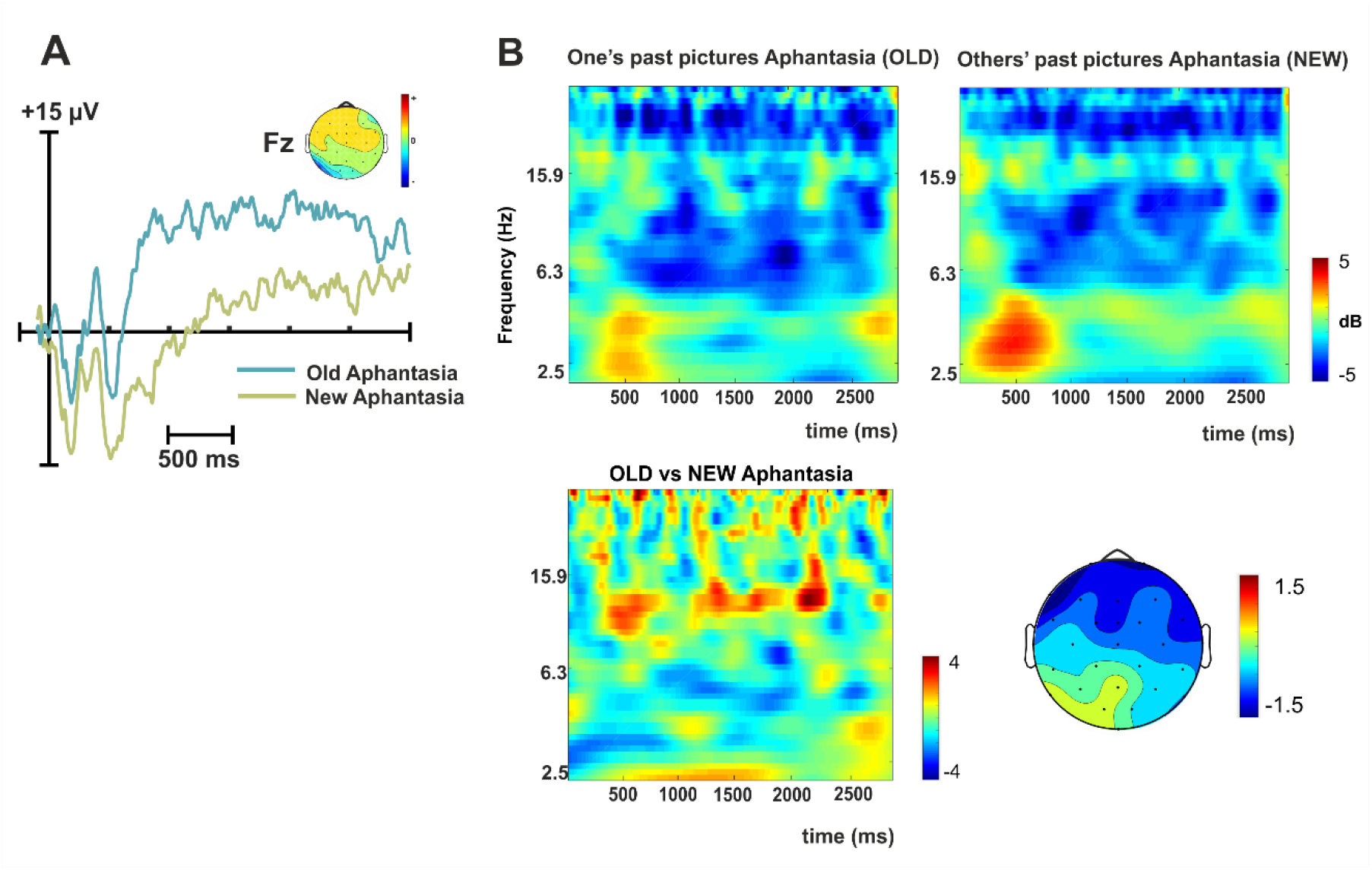
**(A)** Event-related potentials (ERPs) for experiment 3 (case GB) for Old and New conditions at Fz. Old pictures elicited greater ERP amplitudes than did fronto-central scalp regions. **(B)** Changes in spectral power (averaged over all scalp sensors) elicited by pictures related to own personal events (old) compared to others’ events (new). Decreased theta power was observed in old responses compared to fronto-central regions of the scalp.

The time-frequency analysis, however, showed no clear differences between Old and New responses in case GB (Figure 7B). Although this lack of visible theta effect for GB was in contrast to the theta decrease to Old response in participants from experiment 1, Crawford t-test analysis between the two sets of data showed that they were not statistically different (all, p > 0.05).

## Discussion

Real-life experience is characterized by continuous inputs carved by memory systems into an organized, yet intricate, memory network of episodic event representations. How our memory systems are capable of retrieving our unique past episodic experience is a current hallmark in the Autobiographical Memory (AM) research field. In the current study, we proposed a novel methodological approach that combined the use of a wearable camera and implementation of a deep neural network-based algorithm to assess how individuals retrieve specific event episodes from their own past daily life routines. This approach was successful in automatically identifying pictures sequence that represented unique episodic events, thereby reducing the number of “items” and allowing us to explore memory retrieval for the entire past experience relatively rapidly in a later memory test. Our findings revealed that participants were highly accurate in recognizing pictures depicting their own past but that the quality of the retrieved memories changed with the passing of time. Specifically, we found that participants tended to “Remember” with greater accuracy the memory episodes cued by pictures one week after their encoding (test T1), and that this shifted towards a higher proportion of “Know” responses two weeks after (test T2) and 6 to 14 months after (test FU) encoding, thereby suggesting that forgetting was a consequence of the passing time. In line with this behavioral pattern, we found that two well-known electrophysiological brain markers of successful episodic memory retrieval, the Old/New ERP effect and theta neural oscillations, appear clearly associated with the participants’ ability to correctly recognize pictures that cued self-experienced past events from real life, and less clearly, but also evidently, to neutral pictures depicting indoor and outdoor scenes encoded in the lab. Finally, in an attempt to validate this experimental approach at the single-subject level, we presented the data from a single case with Aphantasia, who showed a lack of visual imagery despite having all other cognitive abilities intact.

In experiment 1, we examined the ability of healthy adults to retrieve autobiographical memories of real-life event episodes encoded over a period of a week. We found that participants were highly accurate in doing so but that the quality of the retrieved memories decreased over time, as they reported that picture cues were less prone to eliciting a vivid recollection of the episodic context associated with the picture. This increase in the feeling of familiarity might be explained by the expected decline in memory strength over time that could cause the loss of contextual features, attributed to an inherent time-dependent memory transformation due to consolidation processes. Indeed, during consolidation, memories could lose specific details, becoming more schematic and context-independent with a resulting ‘semantization’ (Winocur & Moscovitch, 2011). These changes could be structurally related to a reorganization of the representational patterns within the hippocampus (Dandolo & Schwabe, 2018), shifting the participants’ judgments from relying on more detailed information when memories are recent, to more gist-based judgments when these memories are remote.

A widely accepted idea is that the retrieval of AMs engages multiple cognitive processes, including the controlled searching process triggered by the cue to a combination of sensory, emotional, and perceptual elements that are needed to come together and emerge as a specific episodic memory (Conway, 2009). This complex process is thought to engage the coordination of several neural networks including the medial prefrontal cortex (PFC), posterior parietal cortex, and medial temporal lobe (MTL) regions, including the hippocampus (Svoboda & Levine, 2009). Previous studies have also attempted to establish a specific role for each of these brain regions. For example, the involvement of the PFC has been attributed to the need to engage a self-reference process and the parietal regions to strategic search and attentional demands (St. Jacques et al., 2011). Our study was blind to the neural sources that were involved at retrieval but, we did find that successful retrieval of AMs was accompanied by modulations of large-scale neural oscillatory patterns in the theta range (4-8Hz). It has been suggested that, through theta oscillations, the MTL may drive the reciprocal exchange of information with neocortical areas (Sirota et al., 2008). Accordingly, the MTL may actively control the transfer of neocortical information to the MTL itself via theta-phase biasing of neocortical network dynamics (Sirota et al., 2008), as postulated in several computational models of memory (Marr, 1971; McClelland et al. 1995; Rolls, 2000; Treves & Rolls, 1994), and this may also account for recent evidence from human intracortical recordings (Foster et al., 2013) and, noninvasively, via magnetoencephalography (Fuentemilla et al., 2014) of MTL-neocortical theta phase-locking during the retrieval of AMs. An interesting issue that remains elusive is why some studies find theta power increases with successful retrieval while others find the opposite pattern, with theta power decreasing, as in the current study. This issue is a topic of intense debate in the research community (see for example Herweg et al., 2019). Nevertheless, our finding, that theta power engagement tends to decrease as a function of memory age (e.g., T1 vs FU), lends support to the notion that MTL-neocortical theta activity decreases over time. This would address the possibility that the MTL regions become less necessary for the retrieval of remote AMs as opposed to recent AMs and therefore support the prediction from system consolidation theory that posits that memories become hippocampus-independent upon their consolidation over time (Diekelmann & Born, 2010).

Interestingly, naturalistic stimuli have received increasing interest in the design of experimental paradigms with videos, short films, and the expansion of virtual reality. Retrieving naturalistic stimuli, such as autobiographical event episodes, requires exploration through a wide search space, the integration of information across multiple time scales, and the activation of self-referential processes and vividness that typically accompany the successful retrieval of ones’ past (Cabeza et al., 2004; Chen & Caplan, 2017; Chow & Rissman, 2017; Gilboa et al., 2004; McDermott et al., 2009; Roediger & McDermott, 2013). Our findings indicate that cueing AMs through pictures recorded from a wearable camera was effective in eliciting this amalgam of cognitive and neural processes, as the magnitude of the behavioral and EEG effects was large and consistent, whereas these effects were minimal or less clear when the same individuals retrieved picture scenes encoded at the lab. Thus, the possibility of studying the retrieval of AMs from real-life experience and at the individual level represents a promising avenue that may have implications at the clinical level, as this can be applied patient by patient, to study memory functioning in daily life routine. Another advantage of our approach is that it may be useful in exploring AM functioning in participants with unusual characteristics that are difficult to treat at the group level in the general population, such as the subject included here with Aphantasia.

Overall, we provide evidence of a novel methodological approach to exploring AMs that is effective in assessing the ability of an individual to retrieve memories from a single event episode that took place in real-life. Our findings help contribute to the emerging body of literature emphasizing the advantages of using naturalistic material rather than artificial lab-based stimuli to explore the cognitive and neural underpinnings of episodic memory. Current findings may be relevant beyond basic research as they may help improve understanding of AM functioning in the clinic at the individual level.

---

**Supplementary Table 1.** Neuropsychological tests and results from GB (Experiment 3)

| COGNITIVE DOMAIN | COGNITIVE PROCESS | TEST | RAW SCORE | NORMATIVE SCORE |
| --- | --- | --- | --- | --- |
| PERCEPTION | Visual Perception | Poppelreuter Overlapping Figures | 40/40 | 100% |
|  | Spatial Organization | Rey Osterrieth Complex Figure-Copy | 32 | SS=9 |
|  | Face Recognition | Famous faces recognition test-Spanish Adaptation | 69/75 | 92% |
| MEMORY | Immediate Visual Memory | Rey Osterrieth Complex Figure - Immediate recall | 27 (type III) | SS=13 |
|  |  | Wechsler Memory Scale (WMS) III - drawings | 102 | SS =17 |
|  | Delayed Visual Memory | Rey Osterrieth Complex Figure - delayed recall | 27 (type III) | SS = 13 |
|  |  | Wechsler Memory Scale (WMS) III - drawings | 72 | SS = 10 |
|  | Visual Recognition | Wechsler Memory Scale (WMS) III - drawings | 47 | SS = 12 |
|  | Immediate Verbal Memory | Wechsler Memory Scale (WMS) III - Short Stories I | 22 | SS = 6 |
|  | Verbal Learning Index | Wechsler Memory Scale (WMS) III - Short Stories I, B2 (learning) | 5 | SS = 11 |
|  |  | Rey Auditory Verbal Learning Task part A1-A5 | 55 | Z= 0.17 |
|  | Delayed Verbal Memory | Wechsler Memory Scale (WMS) III - Short Stories II | 26 | SS = 10 |
|  |  | Rey Auditory Verbal Learning Task (RAVLT) | 2 | Z = -3.4 |
|  | Verbal Learning Curve | Rey Auditory Verbal Learning Task (RAVLT) A1-A2-A3-A4-A5 | 7-11-10-12-15 | (Z) 0.16; 0.5; -0.64; -0.1; 1.21 |
|  | Resistance to interference | Rey Auditory Verbal Learning Task (RAVLT) | 12 | Z= 2.75 |
|  | Verbal Recognition | Wechsler Memory Scale (WMS) III - Short Stories II | 27 | SS = 11 |
|  |  | Rey Auditory Verbal Learning Task - Recognition | 15 | Z = 0.67 |
|  | Auditory memory (music) | Montreal Battery of Evaluation of Amusia (MBAE)- task 1 (scale) | 26/31 | Z= -0.43 |
| **EXECUTIVE FUNCTION** |  | Wechsler Adult Intelligence Scale (WAIS) III - Digit Span (forward and backward) | 20 | SS = 13 |
|  |  | Wechsler Adult Intelligence Scale (WAIS) III – Letters and numbers | 14 | SS = 15 |
|  |  | Rey Osterrieth Complex Figure - Copy | Type III | - |
|  |  | Wechsler Adult Intelligence Scale (WAIS) III - Cubes | 62 | SS = 18 |
|  |  | Wisconsin Card Sorting Task (WSCT) | 9 errors | T = 50 |
|  |  | Tower of London (TOL) total correct responses | 10 | SS =18 |
|  |  | Tower of London (TOL) total number of excessive numbers | 19 | SS = 11 |
|  |  | Semantic (animals) | 26 | SS = 11 |
|  |  | Phonological (letter *p*) | 16 | SS = 10 |
| **INTELLIGENCE ESTIMATES** |  | Wechsler Adult Intelligence Scale (WAIS) III - Vocabulary | 53 | SS = 14 |
|  |  | Wechsler Adult Intelligence Scale (WAIS) III - Similarities | 22 | SS = 12 |
|  |  | Raven Matrices (General Scale) | 54/60 | Normative score |
| **SELF-RATED MEASURES** |  | Vividness of Visual Imagery Questionnaire Revised (VVIQR)- Spanish version | 8/224 | Z= -5.72 |

## References

Bunzeck, N., & Düzel, E. (2006). Absolute Coding of Stimulus Novelty in the Human Substantia Nigra/VTA. Neuron, 51(3), 369–379. https://doi.org/10.1016/j.neuron.2006.06.021

Cabeza, R., Prince, S. E., Daselaar, S. M., Greenberg, D. L., Budde, M., Dolcos, F., … Rubin, D. C. (2004). Brain activity during episodic retrieval of autobiographical and laboratory events: An fMRI study using a novel photo paradigm. Journal of Cognitive Neuroscience, 16(9), 1583–1594. https://doi.org/10.1162/0898929042568578

Chen, Y. Y., & Caplan, J. B. (2017). Rhythmic activity and individual variability in recognition memory: Theta oscillations correlate with performance whereas alpha oscillations correlate with erps. Journal of Cognitive Neuroscience, 29(1), 183–202. https://doi.org/10.1162/jocn_a_01033

Chow, T. E., & Rissman, J. (2017). Neurocognitive mechanisms of real-world autobiographical memory retrieval: Insights from studies using wearable camera technology. Annals of the New York Academy of Sciences, Vol. 1396, pp. 202–221. https://doi.org/10.1111/nyas.13353

Chow, T. E., Westphal, A. J., & Rissman, J. (2018). Multi-voxel pattern classification differentiates personally experienced event memories from secondhand event knowledge. NeuroImage, 176, 110–123. https://doi.org/10.1016/j.neuroimage.2018.04.024

Conway, M. A. (2009). Episodic memories. Neuropsychologia, 47(11), 2305–2313. https://doi.org/10.1016/j.neuropsychologia.2009.02.003

Crawford, J. R., & Howell, D. C. (1998). Comparing an individual’s test score against norms derived from small samples. Clinical Neuropsychologist, 12(4), 482–486. https://doi.org/10.1076/clin.12.4.482.7241

Dandolo, L. C., & Schwabe, L. (2018). Time-dependent memory transformation along the hippocampal anterior-posterior axis. Nature Communications, 9(1), 1–11. https://doi.org/10.1038/s41467-018-03661-7

Delorme, A., & Makeig, S. (2004). EEGLAB: An open source toolbox for analysis of singletrial EEG dynamics including independent component analysis. Journal of Neuroscience Methods, 134(1), 9–21. https://doi.org/10.1016/j.jneumeth.2003.10.009

Diamond, N., & Levine, B. (2020). Linking details to temporal context reinstatement in recall of real-world experiences. https://doi.org/10.31234/OSF.IO/E2UNT

Diekelmann, S., & Born, J. (2010). The memory function of sleep. Nature Reviews Neuroscience, Vol. 11, pp. 114–126. https://doi.org/10.1038/nrn2762

Dimiccoli, M., Bolaños, M., Talavera, E., Aghaei, M., Nikolov, S. G., & Radeva, P. (2015). SR-Clustering: Semantic Regularized Clustering for Egocentric Photo Streams Segmentation. Computer Vision and Image Understanding, 155, 55–69. https://doi.org/10.1016/j.cviu.2016.10.005

Foster, B. L., Kaveh, A., Dastjerdi, M., Miller, K. J., & Parvizi, J. (2013). Human retrosplenial cortex displays transient theta phase locking with medial temporal cortex prior to activation during autobiographical memory retrieval. Journal of Neuroscience, 33(25), 10439–10446. https://doi.org/10.1523/JNEUROSCI.0513-13.2013

Friedman, D., & Johnson, R. A. Y. (2000). Event-Related Potential (ERP) Studies of Memory encoding and retrieval: A selective rewiew. 28(January), 6–28. https://doi.org/10.1002/1097-0029(20001001)51:1<6::AID-JEMT2>3.0.CO;2-R

Fuentemilla, L., Barnes, G. R., Düzel, E., & Levine, B. (2014). Theta oscillations orchestrate medial temporal lobe and neocortex in remembering autobiographical memories. NeuroImage, 85, 730–737. https://doi.org/10.1016/j.neuroimage.2013.08.029

Fuentemilla, Lluís, Palombo, D. J., & Levine, B. (2018). Gamma phase-synchrony in autobiographical memory: Evidence from magnetoencephalography and severely deficient autobiographical memory. Neuropsychologia, 110, 7–13. https://doi.org/10.1016/j.neuropsychologia.2017.08.020

Fuentemilla, Lluís, Penny, W. D., Cashdollar, N., Bunzeck, N., & Düzel, E. (2010). Theta-Coupled Periodic Replay in Working Memory. Current Biology, 20(7), 606–612. https://doi.org/10.1016/j.cub.2010.01.057

Gilboa, A., Winocur, G., Grady, C. L., Hevenor, S. J., & Moscovitch, M. (2004). Remembering our past: Functional neuroanatomy of recollection of recent and very remote personal events. Cerebral Cortex, 14(11), 1214–1225. https://doi.org/10.1093/cercor/bhh082

Greenberg, D. L., & Rubin, D. C. (2003). The neuropsychology of autobiographical memory. Cortex, 39(4–5), 687–728. https://doi.org/10.1016/S0010-9452(08)70860-8

Grossmann, A., & Morlet, J. (1984). Decomposition of Hardy Functions into Square Integrable Wavelets of Constant Shape. SIAM Journal on Mathematical Analysis, 15(4), 723–736. https://doi.org/10.1137/0515056

Herweg, N. A., Solomon, E. A., Kahana, M. J., & Kahana, M. J. (2019). Theta Oscillations in Human Memory. Trends in Cognitive Sciences, 24(3), 208–227. https://doi.org/10.1016/j.tics.2019.12.006

Jeunehomme, O., & D’Argembeau, A. (2018). Event segmentation and the temporal compression of experience in episodic memory. Psychological Research, 84(2), 1–10. https://doi.org/10.1007/s00426-018-1047-y

Jeunehomme, O., & D’Argembeau, A. (2019). The time to remember: Temporal compression and duration judgements in memory for real-life events. Quarterly Journal of Experimental Psychology (2006), 72(4), 930–942. https://doi.org/10.1177/1747021818773082

Keogh, R., & Pearson, J. (2018). The blind mind: No sensory visual imagery in aphantasia. Cortex, 105, 53–60. https://doi.org/10.1016/j.cortex.2017.10.012

LePort, A. K. R., Mattfeld, A. T., Dickinson-Anson, H., Fallon, J. H., Stark, C. E. L., Kruggel, F., … McGaugh, J. L. (2012). Behavioral and neuroanatomical investigation of Highly Superior Autobiographical Memory (HSAM). Neurobiology of Learning and Memory, 98(1), 78–92. https://doi.org/10.1016/j.nlm.2012.05.002

LePort, A. K. R., Stark, S. M., McGaugh, J. L., & Stark, C. E. L. (2017). A cognitive assessment of highly superior autobiographical memory. Memory, 25(2), 276–288. https://doi.org/10.1080/09658211.2016.1160126

Levine, B., Turner, G. R., Tisserand, D., Hevenor, S. J., Graham, S. J., & McIntosh, A. R. (2004). The functional neuroanatomy of episodic and semantic autobiographical remembering: A prospective functional MRI study. Journal of Cognitive Neuroscience, 16(9), 1633–1646. https://doi.org/10.1162/0898929042568587

Maris, E., & Oostenveld, R. (2007). Nonparametric statistical testing of EEG- and MEG-data. Journal of Neuroscience Methods, 164(1), 177–190. https://doi.org/10.1016/j.jneumeth.2007.03.024

Marks, D. F. (1973). Visual imagery differences in the recall of pictures. British Journal of Psychology, 64(1), 17–24. https://doi.org/10.1111/j.2044-8295.1973.tb01322.x

Marr, D. (1971). Simple memory: a theory for archicortex. Philosophical Transactions of the Royal Society of London. Series B, Biological Sciences, 262(841), 23–81. https://doi.org/10.1098/rstb.1971.0078

McClelland, J. L., McNaughton, B. L., & O’Reilly, R. C. (1995). Why there are complementary learning systems in the hippocampus and neocortex: Insights from the successes and failures of connectionist models of learning and memory. Psychological Review, 102(3), 419–457. https://doi.org/10.1037/0033-295X.102.3.419

McDermott, K. B., Szpunar, K. K., & Christ, S. E. (2009). Laboratory-based and autobiographical retrieval tasks differ substantially in their neural substrates. Neuropsychologia, 47(11), 2290–2298. https://doi.org/10.1016/j.neuropsychologia.2008.12.025

Nielson, D. M., Smith, T. A., Sreekumar, V., Dennis, S., & Sederberg, P. B. (2015). Human hippocampus represents space and time during retrieval of real-world memories. Proceedings of the National Academy of Sciences of the United States of America, 112(35), 11078–11083. https://doi.org/10.1073/pnas.1507104112

Nyhus, E., & Curran, T. (2010, June). Functional role of gamma and theta oscillations in episodic memory. Neuroscience and Biobehavioral Reviews, Vol. 34, pp. 1023–1035. https://doi.org/10.1016/j.neubiorev.2009.12.014

Palombo, D. J., Alain, C., Söderlund, H., Khuu, W., & Levine, B. (2015). Severely deficient autobiographical memory (SDAM) in healthy adults: A new mnemonic syndrome. Neuropsychologia, 72, 105–118. https://doi.org/10.1016/j.neuropsychologia.2015.04.012

Palombo, D. J., Sheldon, S., & Levine, B. (2018). Individual Differences in Autobiographical Memory. Trends in Cognitive Sciences, Vol. 22, pp. 583–597. https://doi.org/10.1016/j.tics.2018.04.007

Patihis, L., Frenda, S. J., LePort, A. K. R., Petersen, N., Nichols, R. M., Stark, C. E. L., … Loftus, E. F. (2013). False memories in highly superior autobiographical memory individuals. Proceedings of the National Academy of Sciences of the United States of America, 110(52), 20947–20952. https://doi.org/10.1073/pnas.1314373110

Rissman, J., Chow, T. E., Reggente, N., & Wagner, A. D. (2016). Decoding fmri signatures of real-world autobiographical memory retrieval. Journal of Cognitive Neuroscience, 28(4), 604–620. https://doi.org/10.1162/jocn_a_00920

Roediger, H. L., & McDermott, K. B. (2013). Two types of event memory. Proceedings of the National Academy of Sciences of the United States of America, Vol. 110, pp. 20856–20857. https://doi.org/10.1073/pnas.1321373110

Rolls, E. T. (2000). Memory Systems in the Brain. Annual Review of Psychology, 51(1), 599–630. https://doi.org/10.1146/annurev.psych.51.1.599

Sirota, A., Montgomery, S., Fujisawa, S., Isomura, Y., Zugaro, M., & Buzsáki, G. (2008). Entrainment of Neocortical Neurons and Gamma Oscillations by the Hippocampal Theta Rhythm. Neuron, 60(4), 683–697. https://doi.org/10.1016/j.neuron.2008.09.014

St. Jacques, P. L., Kragel, P. A., & Rubin, D. C. (2011). Dynamic neural networks supporting memory retrieval. NeuroImage, 57(2), 608–616. https://doi.org/10.1016/j.neuroimage.2011.04.039

Svoboda, E., & Levine, B. (2009). The effects of rehearsal on the functional neuroanatomy of episodic autobiographical and semantic remembering: A functional magnetic resonance imaging study. Journal of Neuroscience, 29(10), 3073–3082. https://doi.org/10.1523/JNEUROSCI.3452-08.2009

Treves, A., & Rolls, E. T. (1994). Computational analysis of the role of the hippocampus in memory. Hippocampus, 4(3), 374–391. https://doi.org/10.1002/hipo.450040319

Tulving, E. (2002). Episodic Memory: From Mind to Brain. Annual Review of Psychology, 53(1), 1–25. https://doi.org/10.1146/annurev.psych.53.100901.135114

Wilding, E. L. (1999). Separating retrieval strategies from retrieval success: an event-related potential study of source memory. Neuropsychologia, 37(4), 441–454. https://doi.org/10.1016/s0028-3932(98)00100-6

Winocur, G., & Moscovitch, M. (2011). Memory transformation and systems consolidation. Journal of the International Neuropsychological Society, Vol. 17, pp. 766–780. https://doi.org/10.1017/S1355617711000683

Zacks, J. M., & Swallow, K. M. (2007). Event segmentation. Current Directions in Psychological Science, 16(2), 80–84. https://doi.org/10.1111/j.1467-8721.2007.00480.x

Zeman, A., Dewar, M., & Della Sala, S. (2015). Lives without imagery - Congenital aphantasia. Cortex, Vol. 73, pp. 378–380. https://doi.org/10.1016/j.cortex.2015.05.019

Zeman, A. Z. J., Della Sala, S., Torrens, L. A., Gountouna, V. E., McGonigle, D. J., & Logie, R. H. (2010). Loss of imagery phenomenology with intact visuo-spatial task performance: A case of “blind imagination.” Neuropsychologia, 48(1), 145–155. https://doi.org/10.1016/j.neuropsychologia.2009.08.024

